# Injectable polymer-nanoparticle hydrogel for the sustained intravitreal delivery of bimatoprost

**DOI:** 10.1101/2022.09.08.507193

**Authors:** Emily L. Meany, Roxanne Andaya, Shijia Tang, Catherine M. Kasse, Reina N. Fuji, Abigail K. Grosskopf, Andrea l. d’Aquino, Joshua T. Bartoe, Ryan Ybarra, Amy Shelton, Zachary Pederson, Chloe Hu, Dennis Leung, Karthik Nagapudi, Savita Ubhayakar, Matthew Wright, Chun-Wan Yen, Eric A. Appel

## Abstract

Vision impairment resulting from chronic eye diseases, such as macular degeneration and glaucoma, severely impacts patients’ quality of life and poses an immense global financial burden. Current standard of care for such diseases includes daily eye drops or frequent intravitreal (ITV) injections, which are burdensome treatment modalities resulting in low patient compliance. There remains a growing need for easily administered long-acting delivery technologies for prolonging exposure of ocular therapeutics with each administration. Here, we deploy a supramolecular polymer-nanoparticle (PNP) hydrogel for ITV delivery of the glaucoma drug bimatoprost. PNP hydrogels are shear-thinning and self-healing, key properties for injectability, and enable slow release of molecular cargo in vitreous humor (VH) mimics. An in vivo study in New Zealand white rabbits demonstrated intravitreally injected PNP hydrogels form depots that degrade slowly over time, maintaining detectable levels of bimatoprost in the VH up to eight weeks following injection. Ophthalmic examinations and histopathology identified a mild foreign body response (FBR) to the hydrogel, characterized by rare clusters of foamy macrophages and giant cells associated with minimal, patchy fibroplasia. This work shows that PNP hydrogels exhibit numerous desirable traits for sustained drug delivery and further work will be necessary to optimize tolerability in the eye.

## 1. Introduction

Vision impairment, resulting from eye diseases such as macular degeneration and diabetic macular edema, poses an immense global financial burden and tremendously impacts patients’ quality of life.^[1]^ The World Health Organization projects a steady increase in the prevalence of chronic eye diseases over the next ten years, including a 30% increase from 76 to 95.4 million persons with glaucoma and a 20% increase from 195.6 to 243.3 million persons with age-related macular degeneration.^[2]^ Current standard of care utilizes intravitreal (ITV) administration to treat several ocular diseases and ITV is one of the most effective methods for delivering therapies to the retina. To date, there are various approved ITV biologic therapies, including pegaptanib sodium, ranibizumab, aflibercept, brolucizumab, and more recently faricimab.^[3]^ Yet, despite these robust medical breakthroughs for the management of acquired retinal diseases, patient compliance with repeated ITV injection dramatically falls over time and remains a major obstacle to life-long treatment.^[4]^ Several intervention strategies seek to address this obstacle, including the use of long-acting delivery (LAD) technologies to sustain drug exposure, effectively prolonging efficacy and reducing the frequency of injections.^[5]^ By far, the most advanced LAD technology to successfully address patient compliance is the recently FDA-approved drug, Susvimo™, a refillable port delivery system for ranibizumab.^[6]^ This device is surgically implanted at the pars plana and slowly releases ranibizumab into the vitreous humor (VH) of the eye, with a minimum of 24 weeks between refill exchanges. Other approved LAD technologies for ocular use are predominantly biodegradable implants for sustained immunosuppressive steroid delivery, wherein the steroid itself may mitigate any potential immune response to the delivery vehicle.^[7]^ In the face of continuing growth in populations impacted by ocular diseases, advancement of novel targets for the management of these diseases, and the expanding diversity of drug modalities (i.e., new classes of molecules beyond traditional small molecules) for engaging these targets, the development of injectable LAD technologies for controlled and sustained delivery of ocular therapeutics remains an underserved medical need.

Injectable hydrogels are promising candidates for ocular LAD systems, as these technologies possess numerous unique and desirable features.^[8]^ The high water content and tunable mechanical properties of hydrogels afford exceptional modularity and biocompatibility. Hydrogel systems that employ mild gelation mechanisms, such as supramolecular interactions or ionic, pH, and temperature-triggered interactions, maintain an aqueous environment and promote payload stability, contributing to the versatility of these materials in biologic applications.^[8b, 9]^ Although numerous injectable hydrogel formulations are currently in development as long-acting depots in the eye, none have been approved to date.^[10]^ Barriers to success include a high degree of burst release, poorly matched timescales of drug release and depot degradation (potentially resulting in buildup of depot components), complex manufacturing, and a lack of broad compatibility with various payloads.^[11]^ In addition, demonstration of safety over an extended period of time is necessary since the prolonged presence, or potential accumulation of polymer matrix with repeated dosing, may elicit vitreous haze, foreign body responses (FBRs), retinal toxicities, and an increased risk of visual disturbance.^[7d, 12]^

Our lab has previously developed an injectable hydrogel platform comprising poly(ethylene glycol)-block-poly(lactic acid) nanoparticles (PEG-PLA NPs) and hydrophobically-modified hydroxypropylmethylcellulose (HPMC) polymers.^[13]^ When mixed together, these components form multivalent, noncovalent interactions between the HPMC-C12 polymers and the PEG-PLA NPs leading to formation of robust and tunable polymer-nanoparticle (PNP) hydrogels exhibiting shear-thinning and self-healing properties that enable them to be readily injectable.^[14]^ The gelation process is simple and mild, enabling facile scalability and allowing sensitive cargos like proteins, cells, or small molecules to be readily incorporated.^[13c, 15]^ These materials have previously been shown to be well-tolerated and exhibit excellent biocompatibility when deployed in the intraperitoneal space (in rodents), thoracic cavity (in rodents and sheep), and subcutaneous space (in rodents), suggesting a favorable toxicity profile for ocular applications.^[13d, 13e, 16]^ In this work, we sought to further develop PNP hydrogels as a drug delivery platform in the eye by demonstrating ITV administration for sustained delivery of the glaucoma therapeutic bimatoprost. We sought to evaluate the tolerability of this material platform for the first time in the highly sensitive tissue of the eye. In this work, we characterized the mechanical properties and depot formation of PNP hydrogels in vitro, followed by analysis of the release of bimatoprost from the hydrogel in several in vitro assay designs. We conducted an in vivo study in New Zealand white (NZW) rabbits to evaluate the drug release kinetics and characterize the tolerability profile of these materials over time. We found the PNP hydrogel extended the effective half-life of bimatoprost, maintaining measurable levels of the drug in VH for up to eight weeks, and elicited a mild and localized FBR associated with minimal fibroplasia.

## 2. Results and Discussion

### 2.1. PNP hydrogel for intravitreal extended drug release

In considering biomaterial technologies as LAD systems for ITV drug delivery, several key desirable traits must be targeted: (i) injectability under reasonable forces for a medical practitioner, (ii) rapid self-healing to minimize burst release of entrapped therapeutic cargo, (iii) biocompatibility to ensure minimal FBR in the eye, (iv) similar rates of depot degradation and cargo release to minimize buildup of depot material following repeat injections, and (v) flexibility to accommodate different classes of cargo spanning a diverse array of biologics or small molecules. The hierarchical construction of PNP hydrogels that are self-assembled through dynamic, non-covalent crosslinking interactions between dodecyl-modified HPMC (HPMC-C12) polymers and PEG-PLA NPs, enables facile encapsulation of a broad range of molecular cargos through simple mixing.^[15c]^ The dynamic crosslinking interactions allow facile injection through standard 30-gauge needles typically used for ITV administration.^[17]^ Further, as these materials can be designed such that most molecular cargo is entrapped within the hydrogel network, cargo release can be controlled by the slow erosion of the hydrogel material over time, making the timescales of cargo release and material degradation similar, mitigating potential buildup of hydrogel components.

In this work, we developed PNP hydrogels for the encapsulation and sustained release of bimatoprost following ITV administration, wherein the hydrogel would release drug as it degrades over time in the VH (**Figure 1**). Bimatoprost is a glaucoma medication that is FDA-approved for use in an intracameral implant and was selected for this work owing to its demonstrated ocular tolerability up to 20 μg from Phase I/II clinical trials.^[18]^ We first sought to investigate a PNP hydrogel formulation comprising 2 wt% HPMC-C12 and 10 wt% PEG-PLA NPs, denoted PNP-2-10, based on prior work showing this formulation enables the slowest release rate for various molecular cargo.^[13d, 19]^ We loaded these materials with bimatoprost and evaluated the drug-loaded materials both in vitro and in vivo in New Zealand white rabbits.

**Figure 1.**
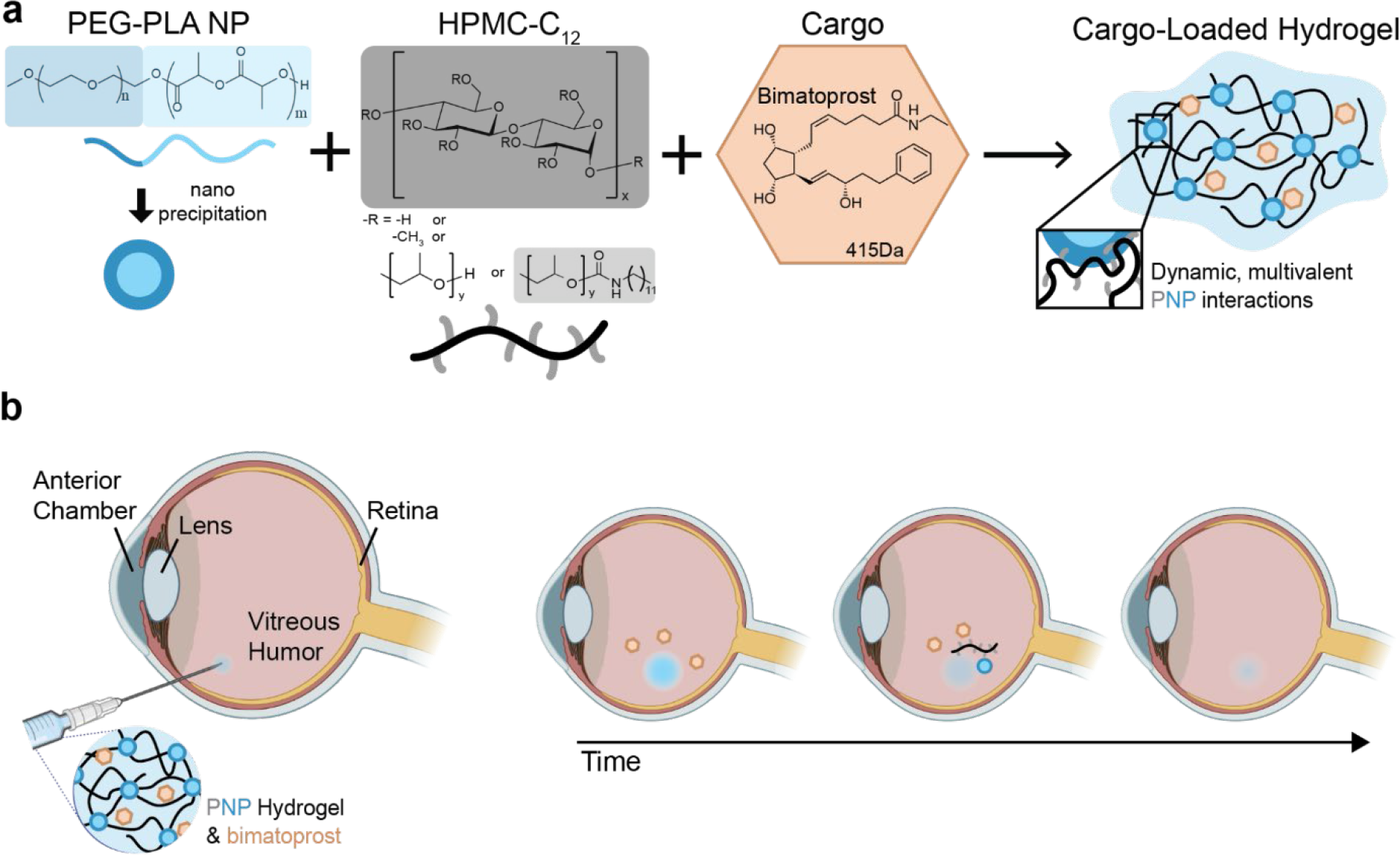
Schematic of the preparation of bimatoprost-loaded polymer-nanoparticle (PNP) hydrogels and their application for prolonged drug release in the vitreous humor (VH) by intravitreal (ITV) injection. **a**. When solutions of PEG-PLA NPs, HPMC-C12, and bimatoprost are mixed, cargo-loaded PNP hydrogels are formed. **b**. PNP hydrogels loaded with bimatoprost are shear-thinning and self-healing, enabling ITV injection and formation of a sustained delivery depot for extended release of bimatoprost in the VH.

### 2.2 Hydrogel development and characterization

PNP hydrogels are facile to make via gentle syringe mixing, where one syringe is loaded with a stock solution of HPMC-C12 and the other is loaded with a stock solution of PEG-PLA NPs and drug cargo (**Figure 2a**). This process of mixing produces a homogenous hydrogel pre-prepared in a syringe for injection. We characterized the rheological properties of PNP hydrogels prepared with and without bimatoprost to ensure the drug cargo does not interfere with the critical properties for injectability and depot formation (**Figure 2b-e**).^[17, 20]^ Frequency-dependent oscillatory shear rheology showed that the frequency responses of the materials were unchanged with encapsulation of bimatoprost. Indeed, both formulations showed robust solid-like properties across the entire range of timescales evaluated, evident from the storage modulus G’ being larger than the loss modulus G’’ (**Figure 2b**). Stress-controlled flow sweep measurements revealed that both materials exhibit static yield stresses of approximately ~500 Pa (**Figure 2c**), and steady shear flow rheology demonstrated robust shear-thinning behavior with dramatically reduced viscosities observed at high shear rates (**Figure 2d**). These characteristics are indicative of a material that will yield and shear-thin, enabling injection through a syringe and needle.^[17]^ It is also important that this material recovers it’s mechanical properties rapidly following injection to limit drug burst release and form a robust depot for extended release.^[20]^ To assess the self-healing of our materials, we applied consecutive periods of high (10 rad s^−1^) and low (1 rad s^−1^) stress to the hydrogel formulations and observed drops in viscosity at high stress and rapid recovery of the material properties at low stress over several cycles, indicating the high stress disrupts the non-covalent PNP interactions responsible for crosslinking in these systems, and these interactions reform once the stress is removed (**Figure 2e**). To further evaluate injectability of the PNP gels, we fit the steady shear flow rheology data with the power law 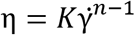, which relates viscosity (η) and shear rate 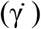 for non-Newtonian complex fluids, and extracted values for the consistency index (*K*) and shear-thinning parameter (*n*).^[17]^ These values fall within the region of “injectability” on an Ashby-style plot of *K* and *n* previously demonstrated to capture the injection forces that are applicable by healthcare professionals (**Figure 2f**).^[17]^ Further, we demonstrated that both empty and bimatoprost-loaded PNP hydrogels were easily injected through a 30-gauge needle, and rapidly formed a robust depot in a VH mimic (**Figure 2g**).

**Figure 2.**
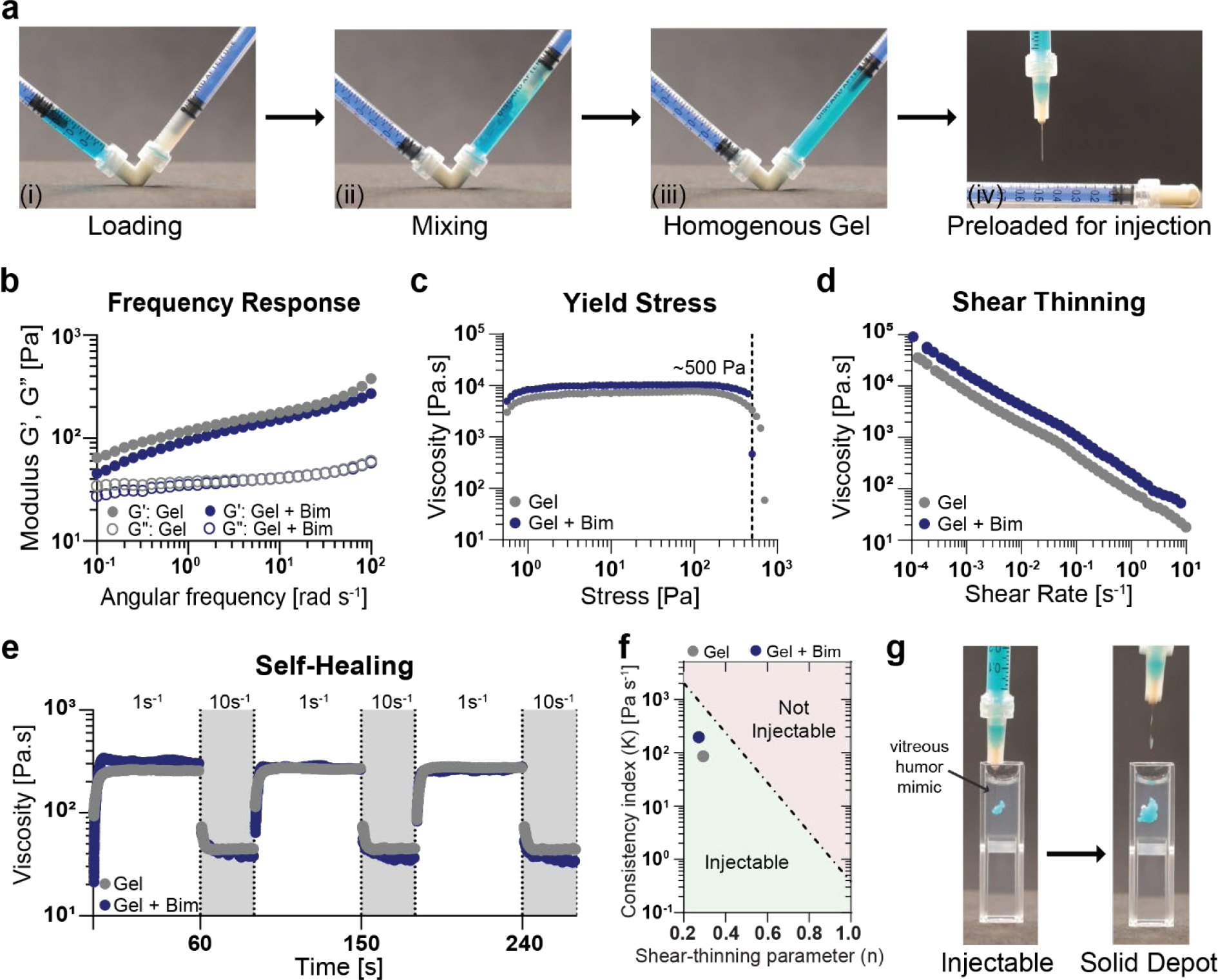
Drug-loaded PNP hydrogels are simple to prepare and their mechanical properties, unimpacted by bimatoprost incorporation, enable injection and depot formation in VH mimic. **a**. PNP hydrogels are prepared by first (i) loading PEG-PLA NPs and cargo (blue dye) into one syringe (left) and HPMC-C_12_ into another (right), (ii) fitting an elbow mixer and mixing back and forth to yield (iii) a homogeneous gel that is (iv) pre-loaded into a syringe for injection. **b**. Oscillatory shear rheology of PNP-2-10 hydrogels (e.g., 2wt% HPMC-C12 + 10wt% PEG-PLA NPs) with (navy blue) or without (gray) bimatoprost (0.25 mg mL^−1^) shows G’ (storage modulus) dominates over G’’ (loss modulus), indicating robust solid-like properties that are not impacted by drug cargo. **c**. Low-to-high shear rheology revealing robust yield stress behavior. **d**. High-to-low shear rheology revealing shear-thinning behavior. **e**. Step-shear rheology measurements demonstrating that PNP hydrogel viscosity drops under high shear (10 s^−1^) and recovers rapidly at low shear (1 s^−1^) over repeated cycles. **f**. Analysis of the flow sweep behavior yields consistency index (*K*) and shear-thinning parameter (*n*) that indicate PNP hydrogels fall into the domain of injectability by healthcare professionals (*K* = 195.5 and 85.62 Pa s^−1^ and *n* = 0.27 and 0.29 for hydrogels prepared with and without bimatoprost, respectively). **g**. PNP hydrogel (50 μL) is easily injected through a 30-gauge needle into VH mimic and self-heals to form a solid depot.

While these PNP hydrogels have been used for controlled delivery in various tissues (e.g., peritoneal cavity, thoracic cavity, and subcutaneous tissue) in multiple animal models (e.g., mice, rats, and sheep), this work is the first to pursue ITV administration in the sensitive environment of the eye. ^[13d, 13e, 16, 21]^ To better visualize and understand how the PNP hydrogels would behave in the VH of the eye, we first analyzed their depot formation and model cargo release in a VH mimic composed of agar and hyaluronic acid (**Figure 3**).^[22]^ For comparison, we evaluated the administration of PNP hydrogels containing either fluorescein (a model small molecule similar in size and hydrophobicity to bimatoprost) and albumin-FITC (a model protein) through a 30-gauge needle into a cuvette containing VH mimic. Bolus administrations of PBS formulations of each dye were used as controls to assess the prolonged delivery of these model molecules from the hydrogel depots (**Figure 3a, c**). The cargo-loaded PNP hydrogels immediately formed depots locally retaining the cargo, whereas cargo administered in PBS boluses rapidly diffused from the injection site following the path of the needle. Over the course of the release experiment, the PNP hydrogel slowed release of both model cargos from the injection site, increasing the effective half-life of retention of fluorescein by over 7.5-fold and of albumin-FITC by over 3.7-fold compared to the control PBS bolus (**Figure 3b, d**). We also observed minimal swelling of the PNP hydrogels over time, and the depot remained suspended, not sinking or floating in the VH mimic, indicating that these materials should remain suspended when injected into the vitreous chamber of the eye rather than contacting the sensitive tissues (e.g. lens, ciliary body, or retina) of the eye.

**Figure 3.**
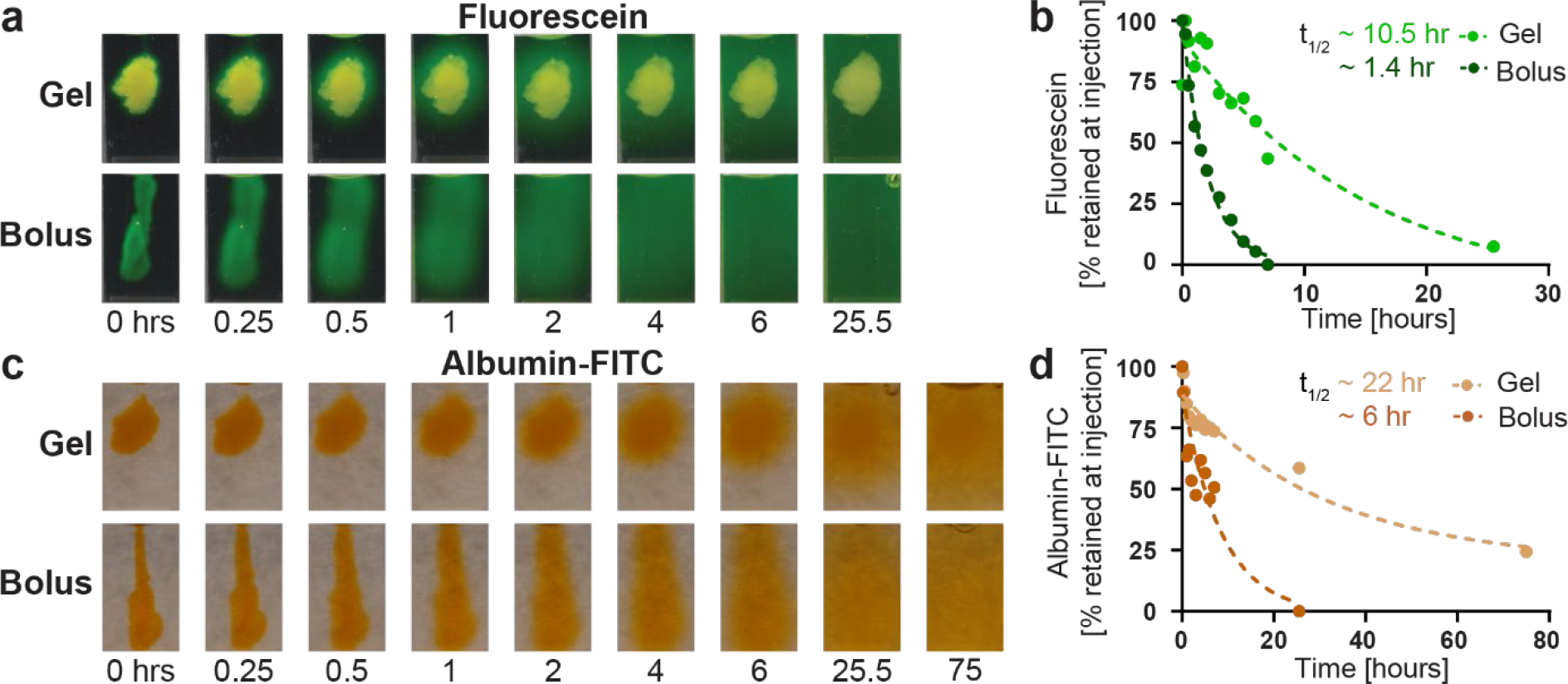
PNP hydrogel slows the release kinetics of model cargo in VH mimic compared with a bolus of model cargo prepared in PBS. PNP hydrogels or PBS (50 μL) containing either **a/b**. fluorescein (0.25 mg mL^−1^) or **c/d**. albumin-FITC (10 mg mL^−1^) was injected into a cuvette containing a VH mimic and imaged over time. Release of the model cargo was quantified using ImageJ software and fit with a one-phase exponential decay in GraphPad Prism to determine a half-life of release for each compound from each vehicle (PNP hydrogel or PBS bolus).

To characterize the release behavior of bimatoprost from the PNP hydrogels, we conducted infinite sink release assays in three separate configurations to later identify which release study formats better recapitulate physiological conditions. We implemented two static assays (**Figure 4a**) and one dynamic release assay (**Figure 4c**). The first static release assay employed a glass capillary tube, where the hydrogel was injected into the bottom of the tube and confined with a single surface in contact with release buffer (PBS). The release buffer was removed and replaced at each time point to maintain infinite sink conditions and to evaluate bimatoprost release over time. We hypothesized this configuration would mimic hydrogel injected into a treatment area surrounded with little biological fluid and minimal tissue motion, such as the subcutaneous space, and have previously demonstrated good correlation between in vitro and in vivo release characteristics.^[20, 23]^ The second assay employed a static dialysis release setup, whereby hydrogel was injected into a microcentrifuge tube with dialysis membrane walls and placed in a sealed tube containing release buffer (PBS). At each time point, a small proportion of the release buffer was removed (~5% total volume) for bimatoprost quantification and replaced with fresh buffer. We hypothesize this configuration would better mimic hydrogel administered into an area with extensive exposure to fluid without significant motion, such as VH.

**Figure 4.**
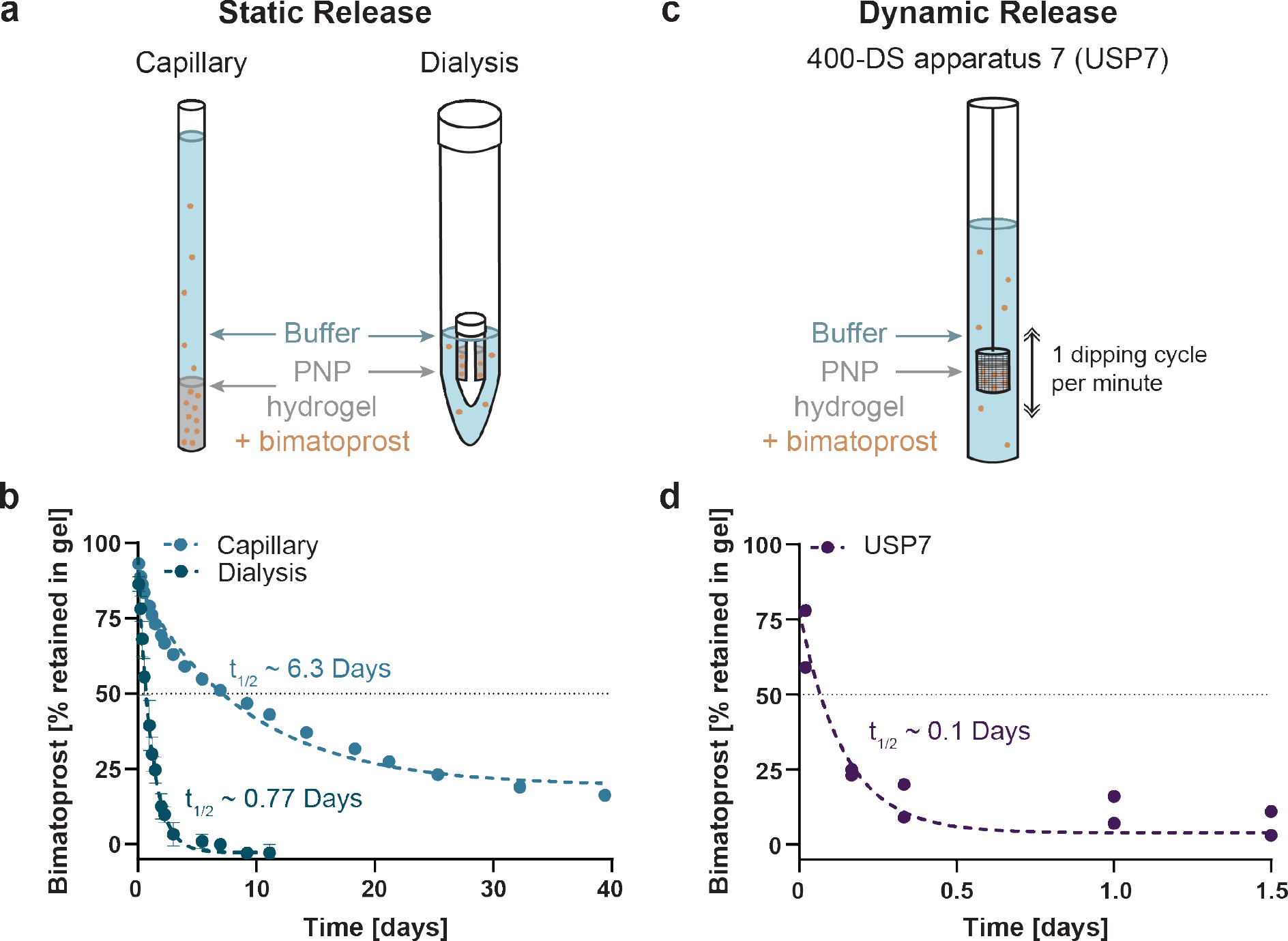
In vitro release of bimatoprost from PNP hydrogel. **a**. Schematic of two static release assays wherein 100 μL of PNP hydrogel with bimatoprost cargo (0.25 mg mL^−1^) was injected into the bottom of a capillary or dialysis tube and PBS buffer added. Buffer was sampled over time to quantify bimatoprost release. **b**. Quantification of bimatoprost release over time (n = 3) as determined by LC-MS. Data are shown as mean ± SD and fit with a one-phase decay in GraphPad Prism and half-lives calculated. **c**. Schematic of dynamic release assay with 400-DS apparatus 7 wherein 100 μL of PNP hydrogel with bimatoprost (0.74 mg mL^−1^) was injected into the mesh basket and PBS buffer added. The basket was dipped at 1 cycle per minute and buffer sampled over time. **d**. Quantification of bimatoprost release over time (n = 2) as determined by RP-CAD. Data are shown as individual points and fit with a one-phase exponential decay in GraphPad Prism and half-life calculated.

Release profiles from both static assays showed no evidence of burst release, which is a key advantage of the PNP hydrogel over other injectable hydrogel systems.^[8b]^ The capillary configuration yielded a half-life of release of 6.3 days, which was much longer than the half-life observed in the dialysis configuration, which was only 0.77 days (**Figure 4b**). This observation suggests that extensive fluid exposure can lead to faster-than-expected erosion of the PNP hydrogels. Cargo release from polymeric materials can be described by the exponential relationship derived by Ritger and Peppas: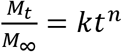 where the release constant (*k*) and diffusional exponent (*n*) are characteristic of the release mechanism.^[24]^ Pure Fickian diffusion yields an *n* value of 0.43 for a sphere, whereas values above this indicate some degree of anomalous release on account of, for example, swelling, while values below this indicated sub-diffusive release. Fitting of the release curves for both static release assays yielded diffusional exponents of *n* ~ 0.39 for the capillary setup and *n* ~ 0.66 for dialysis setup (**Figure S1**). These results may indicate that excess buffer and fluid motion (e.g, observed in the floating dialysis cassette), which doesn’t fully recapitulate the gelatinous mixture of hyaluronic acid and other biopolymers, proteins, and cells of the VH, leads to anomalous release in the dialysis tube, whereas sub-diffusive release is observed when erosion is limited in the capillaries. ^[25]^

To evaluate the release of bimatoprost from the PNP hydrogels in a dynamic assay, we used an Agilent 400-DS apparatus (USP7), often used for dissolution testing, batch analysis, and quality control of drug-eluting stents and medicated contact lenses, among other drug delivery technologies.^[26]^ In these assays, bimatoprost-loaded PNP hydrogels were loaded into a mesh basket that was immersed in release buffer (PBS) and moved up and down with a programmed dipping rate of 1 cycle per minute. The release half-life in this assay was determined to be only 0.1 day, significantly shorter than the release half-lives observed in either static release assay due to the additional shearing forces that lead to the rapid and complete dissolution of the PNP hydrogels. Indeed, the PEG-PLA NP and HPMC-C12 components of the hydrogel were quantified in the releasate and showed similar release profiles as the entrapped bimatoprost (**Figure S2**). This observation suggests that release of the cargo from these hydrogels is driven primarily by dissolution-based erosion of the hydrogels and corroborates the sub-diffusive release observed in the static capillary assays described above. While this USP7 setup does not sufficiently mimic the conditions within VH to capture relevant release timescales, it nevertheless highlights that the PNP hydrogels are physically crosslinked and completely dissolve away on a similar timescale as the release of the encapsulated drug. This feature is crucial as it potentially eliminates the amount of excess material remaining following cargo release, which is a critical attribute for LAD technologies used in the eye. Comparison of these three assays highlights the importance of considering the final application of an LAD material in vivo when selecting a configuration setup for cargo release assays.

### 2.3 In vivo depot and pharmacokinetic characterization

We next proceeded to characterize the PNP hydrogel in vivo in NZW rabbits, a well-established preclinical model for investigating the safety of ocular therapeutic candidates.^[27]^ For these studies, we targeted a bimatoprost dose of 8 μg and an injection volume of 50 μL per eye to minimize any potential adverse effects due to the drug cargo or administration protocol and match the dosing used for in vitro characterization. Bilateral ITV injections delivered PNP hydrogels without bimatoprost (Gel) to two rabbits (four eyes total), and PNP hydrogels with bimatoprost (Gel + Bim) to six rabbits (twelve eyes total). Both formulations were confirmed to be endotoxin-free prior to administration. Rabbits were monitored for two months following ITV injection to assess ocular tolerability by ophthalmic examinations (OE) comprised of slit-lamp biomicroscopy, indirect ophthalmoscopy, and rebound tonometry for intraocular pressure (IOP) measurements. Wide-field color fundus and ultrasound imaging were used to monitor the appearance of the hydrogel and spectral domain - optical coherence tomography (SD-OCT) was used to qualitatively assess hydrogel-related changes to retinal layer thickness. Following euthanasia, ocular tissues were collected for pharmacokinetic (PK) analysis of bimatoprost in VH and histopathological analysis to evaluate any microscopic changes (**Figure 5a**).

**Figure 5.**
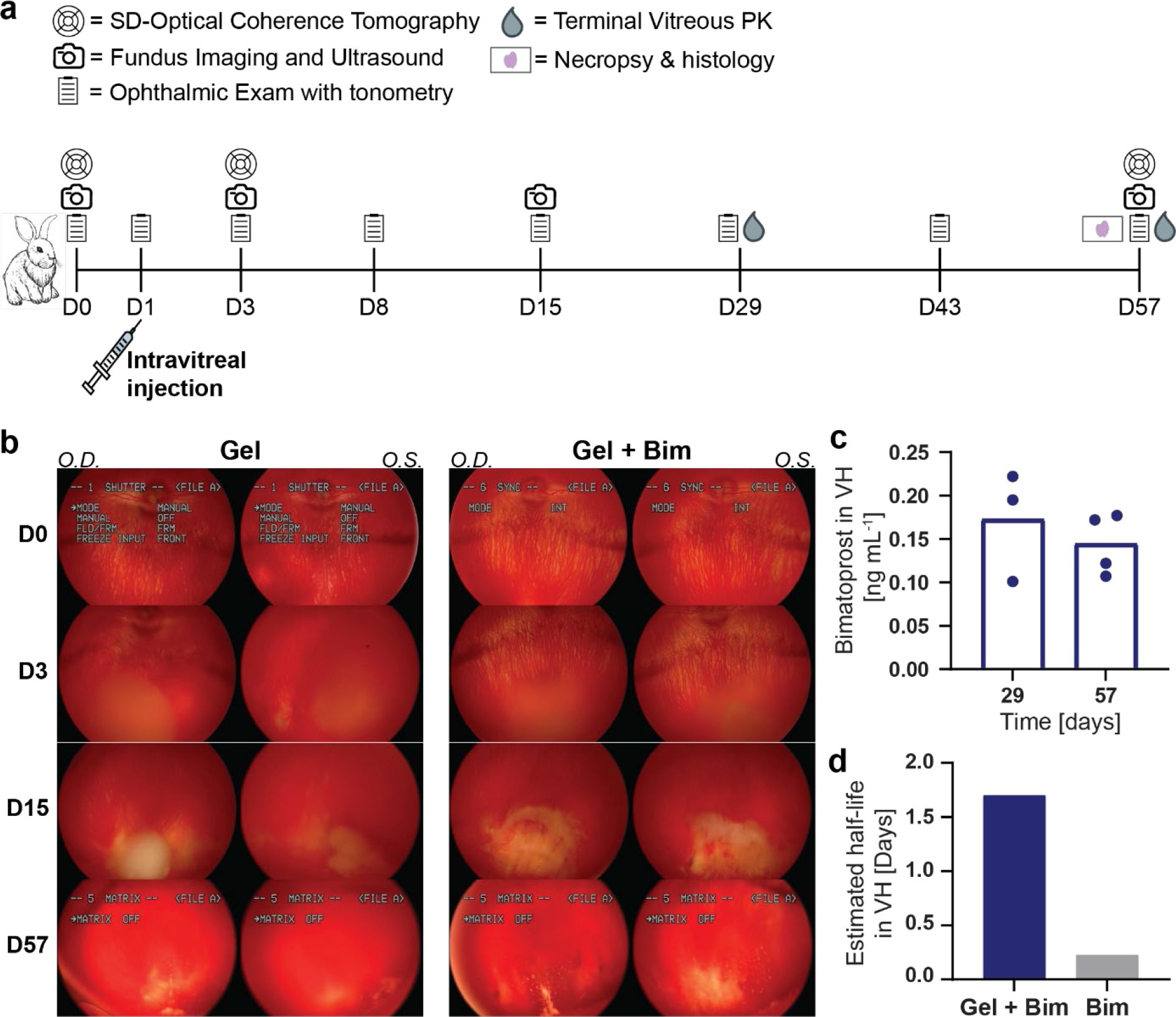
In vivo application of PNP hydrogel in New Zealand white (NZW) rabbit model. **a**. Timeline of rabbit tolerability and PK study. PNP hydrogel alone (Gel) or with bimatoprost (0.16 mg mL^−1^) (Gel + Bim), was injected intravitreally into both eyes of rabbits on day 1 (50 μL injection). Ocular tolerability was evaluated over time by ophthalmic exam (OE) with tonometry. Wide-field color fundus and ultrasound imaging were used to monitor the appearance of the hydrogel and SD-OCT was used to qualitatively assess retinal layer thickness. Globes and VH were collected at terminal time points on days 29 and 57 for histology and PK analysis. **b**. Representative color fundus images show clear depot formation on days 3 and 15 with substantial degradation of the depot on day 57 (*O.D*. = oculus dexter, *O.S*. = oculus sinister). **c**. Bimatoprost in VH was quantified by LC-MS at terminal time points on days 29 (n = 3) and 57 (n = 4). **d**. VH half-life of bimatoprost dosed using PNP hydrogel was calculated using an administered dose of 8μg bimatoprost in an estimated 1.5 mL VH and one-phase decay fit in GraphPad Prism. VH half-life of bimatroprost alone was predicted using a pharmacokinetic model based on its physiochemical parameters.^[30]^

Immediately following administration, the hydrogels were visualized as a smooth, faintly opaque depot in the inferior VH. Few instances of spherical foci, presumed to be air bubbles, were observed at the surface or within the depot in both the Gel and Gel + Bim groups. By day 3, an increase in spherical foci were visualized within the hydrogel in both groups and the edges of the hydrogel depots appeared discontinuous. As the study progressed, the hydrogel depots were subjectively less smooth and defined, with strand-like edges and refractive precipitates or cell-sized particles visible near or within the depot. These precipitates and particles were presumed to be components of the hydrogel undergoing degradation and dissolution and were more numerous in eyes dosed with Gel + Bim. Fundus photographs taken at regular intervals confirmed the hydrogel depot was diminishing in size and definition over time, indicating degradation and dissolution of the hydrogel depots (**Figure 5b**).

At days 29 and 57, two rabbits in the Gel + Bim group were euthanized (n = 4 eyes per timepoint) and VH was collected and analyzed by LC-MS to measure bimatoprost concentration. Detectable levels of bimatoprost were found even at day 57, indicating the PNP hydrogel was successful at sustaining the release of this small molecule. We estimated the effective elimination half-life of bimatoprost from the eye to be 1.7 days using the initial dose of 8 μg in a standard volume range for rabbit VH (1.15 - 1.7 mL).^[28]^ This estimated half-life falls within the range of half-lives observed from the in vitro static release assays discussed above. As the elimination half-life of bimatoprost following IV administration is reported to be only 45 minutes, the effective half-life measured here for the PNP hydrogel based formulations in VH represents an almost two-order-of-magnitude improvement.^[29]^ To the best of our knowledge, the half-life of bimatoprost in VH has not been published, so we used a pharmacokinetic model built on experimental rabbit data and physiochemical properties of various drugs (molecular weight and lipophilicity) to estimate the half-life.^[30]^ When applied to bimatoprost, the model conservatively predicted a half-life of 0.225 days, in agreement with known elimination half-lives of other small molecules in VH.^[30]^ Therefore, PNP hydrogel increased the effective elimination half-life of bimatoprost in VH by more than 7.5-fold from 0.225 days to 1.7 days. While a more thorough analysis of pharmacokinetics with additional timepoints is needed, these initial data are promising and demonstrate measurable bimatoprost in VH for almost two months.

### 2.4 In vivo tolerability evaluation

A key parameter for any long-term drug delivery vehicle is tolerability in the sensitive and immune-privileged tissues of the eye. We evaluated possible adverse effects of the PNP hydrogels through several in-life measurements including OE with tonometry, fundus and ultrasound imaging, and SD-OCT. IOP was transiently elevated above the normal range (10 – 20 mmHg) for both groups (Gel and Gel + Bim) immediately following administration, which is expected for ITV injection (**Figure 6a**). In subsequent intervals, IOP values were found to be below normal, and trace aqueous flare was observed, consistent with ITV injection-related breakdown of the blood-ocular barrier (BOB). While the therapeutic effect of bimatoprost is decreased IOP, a sustained drop in IOP was not expected in these studies as we were not using a disease model exhibiting elevated IOP.^[31]^ As the BOB integrity returned, IOP values returned to normal levels and aqueous flare resolved in a majority of eyes in both groups, and both metrics remained within normal range for the duration of the study. Generally, these IOP findings were more pronounced in eyes dosed with Gel + Bim, presumably on account of bimatoprost’s known impact on BOB integrity that may exacerbate preexisting inflammation.^[32]^

**Figure 6.**
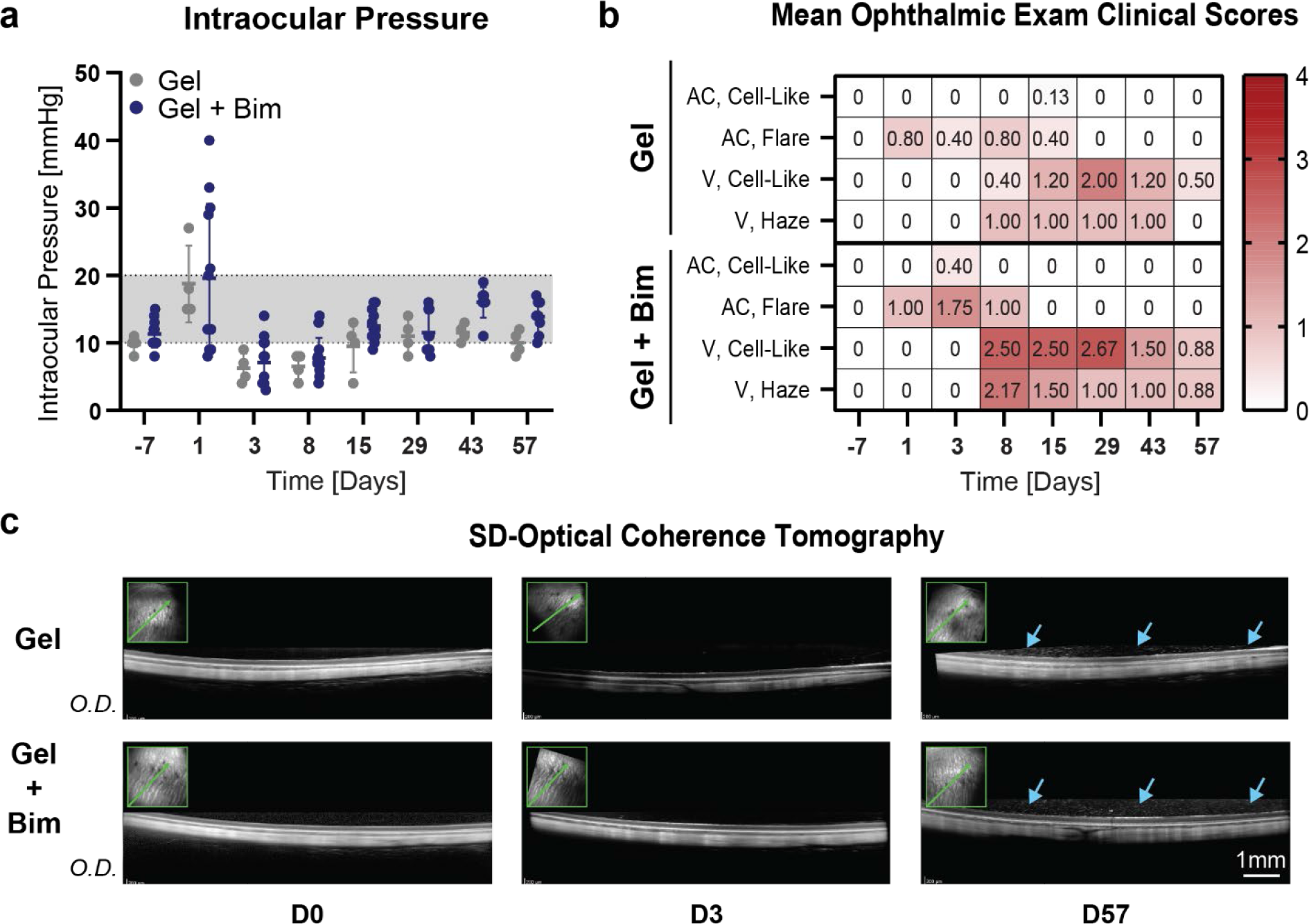
In-life ophthalmic exam (OE) with rebound tonometry and spectral domain – optical coherence tomography (SD-OCT) to assess PNP hydrogel tolerability. **a**. Intraocular pressure (IOP) measurements for eyes injected with PNP hydrogel alone (Gel), or with bimatoprost (0.16 mg mL^−1^) (Gel + Bim), fell within the average range (10 – 20 mmHg). IOP elevation following administration and decrease on days 3 and 8, were expected procedure-related trends following standard ITV injection. Data shown as mean ± SD. **b**. Means of clinical scores for several OE parameters showed presence of cells and cell-like material in the VH that mostly resolved and trended towards recovery at the end of the study (*AC* = aqueous chamber, *V* = vitreous). **c**. Representative SD-OCT images demonstrate subjectively no changes in retinal thickness over time but identified an accumulation of hyperreflective debris within the posterior VH (blue arrows) at the terminal timepoint, presumably a product of PNP hydrogel breakdown (*O.D*. = oculus dexter).

As the IOP and aqueous flare associated with injection and bimatoprost subsided, a minor inflammatory reaction was observed in the vitreous. This reaction was characterized by trace to moderate haze and refractive, cell-like material that was likely a mix of inflammatory cells and hydrogel degradant products, suggestive of an FBR to the PNP hydrogel (**Figure 6b**). Qualitatively, the measure of cell-like material peaked at day 29 and resolved by the terminal time point, with slightly greater severity maintained in eyes dosed with Gel + Bim. SD-OCT and ultrasound did not reveal any alterations to the retina, but an accumulation of hyperreflective debris within the VH immediately adjacent to the retinal surface, presumed to be hydrogel degradant products, was apparent in both groups at study termination (**Figure 6c**, **Figure S3**).

Microscopic examination of all eyes was conducted at study termination to identify any changes related to ITV injected PNP hydrogel. Slides were prepared from paraffin-embedded eyes and stained with hematoxylin and eosin (H&E). Minimal to mild infiltration and accumulation of foamy macrophages and multinucleated giant cells in the VH was observed in eyes from both groups (**Figure 7**). The cells were present in clusters anteriorly and inferiorly near the ciliary body (region of hydrogel deposition), scattered in smaller clusters extending inferiorly along the retina toward the back of the eye, and in the iridocorneal drainage angle, likely in the process of clearing from the eye. These cells were also observed in close association with minimal, patchy fibroplasia. The presence of macrophages, and particularly multinucleated giant cells, suggests a mild FBR to the PNP hydrogel. Injection procedure-related minimal to moderate mixed cell inflammation of the bulbar conjunctiva and/or sclera was also observed.

**Figure 7.**
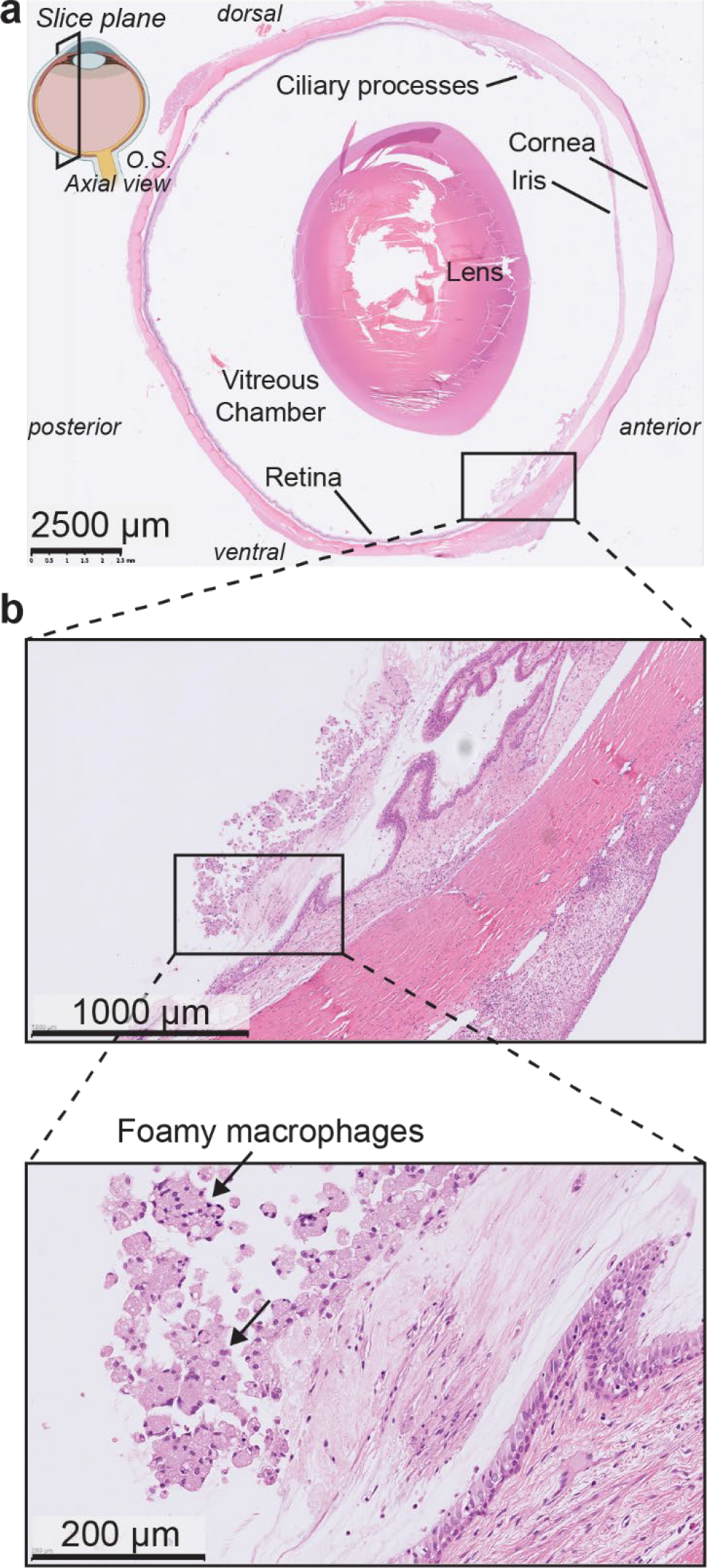
Histopathology of rabbit eyes following ITV administration of PNP hydrogels. Minimal to mild FBR in the VH and along the ventral retina was observed microscopically in response to PNP hydrogel. **a**. Representative image of H&E staining of the full rabbit eye (scale bar = 2500 μm) shows an affected region extending from the ciliary body and along the ventral peripheral retina **b**. Higher magnification of the affected regions shows foamy macrophages.

FBRs have been observed in rabbits following ITV injection of other materials, with FBR-associated inflammation observable during OE soon following injection or implantation.^[33]^ One contributing factor to the observed FBR to the PNP hydrogels may be the difference in mechanical properties between the hydrogels and VH. While the PNP hydrogels exhibit low moduli, their storage modulus of ~150 Pa is still higher than the modulus of VH, which has been shown to be between 1 and 10 Pa.^[22, 34]^ In contrast to other LAD technologies in rabbits, OEs captured only a mild inflammatory reaction that subsided over time in this study.^[7d, 33]^ Furthermore, no major retinal changes were observed upon OCT or ultrasound. Despite these favorable observations, microscopic findings of fibroplasia in response to the presence of PNP hydrogels, while relatively mild and patchy, demonstrates that complete tolerability of hydrogel materials in VH remains a significant challenge as such microscopic changes can neither be monitored nor managed in the clinic.

## 3. Conclusion

Populations suffering from chronic eye diseases are projected to increase over the next ten years and current treatment strategies requiring repeated ITV injections are hindered by low patient compliance, highlighting the need for LAD technologies. In this work, we demonstrated first-in-rabbit and first-in-eye use of a supramolecular polymer-nanoparticle (PNP) hydrogel LAD system. These PNP hydrogels exhibit mechanical properties that are challenging to engineer, yet crucial for ITV administration, including injectibility through 30-gauge needles typically employed for ITV injection, as well as rapid self-healing to form a robust solid depot in the eye. We showed PNP hydrogels slowed the release of both a model small molecule and biologic cargo in VH mimic. Using three separate in vitro release assays we evaluated the release behavior of the glaucoma drug bimatoprost, which provided critical insights into how assay parameters influence release kinetics. Bimatoprost exhibited sub-diffusive release with a half-life of 6.3 days from the PNP hydrogels in a static capillary release assay, but anomalous release and a half-life of 0.77 days when surrounded with buffer in a dialysis assay. The physically crosslinked nature of the PNP hydrogels also led to complete dissolution and release of bimatoprost with a half-life of only 0.1 days when exposed to shear forces in a dynamic release assay using a USP7 apparatus. These studies highlight the importance of careful consideration of the target in vivo environment when designing in vitro release studies to best recapitulate what a LAD will experience upon administration in vivo.

Studies of both empty and bimatoprost-loaded PNP hydrogels following ITV administration in NZW rabbits revealed that these materials enable prolonged exposure to the drug. Fundus imaging and ultrasound imaging visualized PNP hydrogel morphology as it degraded over time in VH. Terminal SD-OCT showed no retinal changes, but identified an accumulation of hyperreflective debris, presumed to be hydrogel degradation products, above the retinal surface. OEs and histopathology revealed mild inflammation and FBR to the PNP hydrogel, characterized by infiltration of foamy macrophages and multinucleated giant cells, along with co-localized fibroplasia. While PNP hydrogels were observed to induce a mild FBR, improving upon previously reported LAD systems evaluated in the eye, the resulting patchy fibroplasia poses clinical risks, such as retinal detachment. These studies highlight the ongoing challenges for developing LAD technologies that are well-tolerated in the eye, even when they exhibit biocompatibility in other parts of the body. As the chemical identity, molecular weight and modifications of polymers can all impact their tolerability, future work will focus on optimizing the PNP hydrogel platform to enhance tolerability in the eye and improve suitablilty for ocular drug delivery.

## 4. Experimental Methods

### Materials

Hypromellose (HPMC, meets USP testing specifications), N-methyl-2-pyrrolidone (NMP), 1-dodecylisocynate, N,N-diisopropylethylamine (Hunig’s base), acetone, monomethoxy-PEG (5 kDa), diazobicylcoundecene (DBU), acetic acid, formic acid, diethyl ether, hexanes, dimethyl sulfoxide (DMSO), acetonitrile, albumin-FITC, agar, and fluorescein were purchased from Sigma-Aldrich and used as received. Dichloromethane (DCM) was purchased from Sigma-Aldrich and further dried via cryo distillation. Lactide (LA) was purchased from Sigma-Aldrich and purified by recrystallization in ethyl acetate with sodium sulfate. Sodium hyaluronate (HA, research grade, 1.0-1.8 MDa) was purchased from Lifecore Biomedical. Acetonitrile (HPLC grade; J.T. Baker) and milliQ water were used for all HPLC analysis. Bimatoprost was purchased from Toronto research chemicals.

### HPMC-C_12_ synthesis

HPMC–C_12_ was prepared according to previously reported procedures.^[35]^ HPMC (1.0 g) (SEC MALS: Mw (Đ) = 372.4 kDa (1.43), method previously reported) was dissolved in NMP (40 mL) at room temperature with stirring. Once the polymer had completely dissolved, the reaction was brought to 80 °C and a solution of 1-dodecylisocynate (0.5 mmol) in NMP (5 mL) was added dropwise, followed by Hunig’s base (catalyst, ≈ 10 drops). The reaction was removed from heat and allowed to react with stirring at room temperature for 16 h. The solution was then precipitated from acetone and hydrophobically-modified HPMC was recovered by dialysis against MilliQ water for 3-4 days (MWCO 3.5 kDa) and lyophilization, yielding HPMC-C12 as a white amorphous powder. The polymer was reconstituted as a 60 mg mL^−1^ solution with sterile PBS, pH 7.4, prior to use in hydrogels.

### PEG-PLA synthesis

PEG-PLA was prepared and analyzed as previously reported.^[35]^ Recrystallized LA (10 g) was dissolved in cryo-distilled DCM (50 mL) under nitrogen with mild heating. Methoxy poly(ethylene glycol) (5 kDa; 2.5 g) was heated to 90 °C under vacuum for 30 minutes, allowed to cool slightly under nitrogen, and then dissolved in cryo-distilled DCM (5 mL) with distilled DBU (75 μL; 0.5 mmol; 0.7 mol% relative to LA). The PEG/DBU solution was added rapidly to the LA solution and allowed to stir for 8 min. The reaction mixture was quenched with acetone (500 μL) and acetic acid (~2 drops) and precipitated from excess 50:50 mixture ethyl ether and hexanes. The PEG-PLA copolymer was collected and dried under vacuum to yield a white amorphous powder. DMF GPC: Mw (Đ) = 22.5 kDa (1.07), method previously reported.^[35]^

### PEG-PLA nanoparticle (NP) preparation

NPs were prepared and analyzed as previously reported.^[35]^ A solution (1 mL) of PEG-PLA in 25:75 DMSO:Acetonitrile (50 mg mL^− 1^) was added dropwise to water (10 mL) at a stir rate of 600 rpm. NPs were purified by ultracentrifugation over a filter (MWCO 10 kDa; Millipore Amicon Ultra-15) followed by resuspension in PBS to a final concentration of 200 mg ml^− 1^. NP size and dispersity were characterized by DLS (Wyatt DynaPro PlateReader-II; average diameter = 31.8 nm, PDI = 0.04)

### PNP hydrogel dose formulation

PNP-2-10 hydrogels were formulated with final concentrations of 2 wt% HPMC-C12 and 10 wt% PEG-PLA NPs. HPMC-C12 was dissolved in PBS at 6 wt% and loaded into a luer-lock syringe (1 or 3 mL depending on volume of gel needed). A 20 wt% solution of NPs in PBS was diluted with additional PBS, or PBS containing bimatoprost at the desired concentration, and loaded into a separate luer-lock syringe (1 or 3 mL). The nanoparticle syringe was then connected to a female-female luer-lock elbow and the solution was moved into the elbow until visible at the other end. The HPMC-C12 syringe was then attached to the other end of the elbow with care to avoid air at the interface of HPMC-C12 and the NP solution. The two solutions were mixed for 1 minute or until a homogenous PNP hydrogel was formed. After mixing, the elbow was removed and a needle of the appropriate gauge was attached.

For animal dosing, hydrogel material (~200 μL) with or without bimatoprost (0.16 mg mL^−1^) was stored at 4 °C in sterile syringes enclosed in sterile 50 mL falcon tubes (one per dose) until the day of dosing. On the day of dosing, > 50 μL of material was back-loaded into individual dosing syringes (0.3 mL insulin syringe, 31-gauge x 8 mm needle, one per eye) and the syringe plunger was used to slowly concentrate the gel at the tip of the needle, careful to avoid bubbles in the hydrogel. 50 μL was injected intravitreally per eye.

### PNP hydrogel rheological characterization

Rheological characterization was performed on PNP hydrogels with or without bimatoprost (0.25 mg mL^− 1^) using a TA Instruments DHR-2 stress-controlled rheometer. All experiments were performed using a 20 mm diameter serrated plate geometry at 25 °C with a 500 μm gap. Frequency sweep measurements were performed at a constant 1% strain in the linear viscoelastic regime. Stress sweeps were performed from low to high with steady state sensing and yield stress values extracted. Flow sweeps were performed from high to low shear rates. Step shear experiments were performed by alternating between a low shear rate (0.1 s^− 1^; 60 s) and a high shear rate (10 s^− 1^; 30 s) for three cycles.

### Injectability calculation and Ashby-style plot

High shear data (shear rates 0.1 – 10 rad s^−1^) from flow sweep rheology was fit with the power law 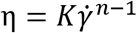 relating viscosity (η) and shear rate 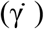 in GraphPad Prism and values for consistency index (*K*) and shear-thinning parameter (*n*) extracted. Values are plotted as points (*n, K*) along with the line defining injectability as previously reported.^[17]^ Injectability is defined as the region with 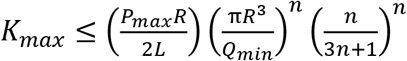 where *P_max_* is the maximum pressure to be exerted during injection, *R* is the diameter of the needle, *L* is the needle length, and *Q_min_* is the minimum desired flow rate. Parameters used: *P_max_* = 2.6 MPa, *R* = 133 μm, *L* = 8.5 mm, and *Q_min_* = 6 mL min^−1^ (50 μL in 0.5 seconds).

### Vitreous humor mimic release assay

50 μL of PNP hydrogel or PBS loaded with either 0.25 mg mL^−1^ fluorescein or 10 mg mL^−1^ albumin-FITC (MW ~ 66 kDa, mol FITC:mol albumin = 14) was injected through a 30-gauge needle into a cuvette containing a VH mimic. Cuvettes were imaged at 0, 0.25, 0.5, 1, 1.5, 2, 3, 4, 5, 6, 7, 25.5, and 75 hrs with dark and light backgrounds for visualization of the dye and gel depot. Dye release was quantified using ImageJ software. Images were separated into red, blue, and green channels and the intensity in the green channel within a region inside the hydrogel depot or initial PBS injection site was measured for each time point, denoted 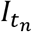. A region in the lower right of the cuvette (outside of the injection site) was used for background intensity measurements to determine when each PBS sample had completely diffused, denoted *t_plateau_*. The fraction of dye remaining in the gel depot at the final time point was determined by comparing the change in background intensity from times *t_0_* to *t_plateau_* for PBS and gel samples: 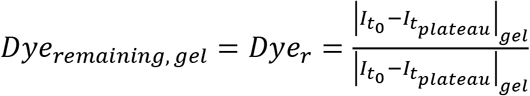. The percent of dye retained in the gel or PBS injection site at each time point *t_n_* was determined by normalizing the intensity as 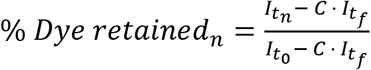, where *t_f_* is the final time point measured and *C* is a constant defined by 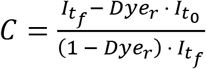. Note, for these samples, *t_f_* = *t_plateau_*.

### In vitro static capillary and dialysis release assays

For capillary release, glass capillary tubes were sealed on one end with epoxy and allowed to cure for at least 24 hours. PNP hydrogel loaded with bimatoprost (100 μL, 0.25 mg mL^−1^) was injected into the bottom of each tube (n = 3) and PBS (400 μL) was injected on top carefully to not disrupt the gel surface. Tubes were sealed with parafilm and stored upright at 37 °C. At each time point, all 400 μL PBS was carefully removed from the tube and replaced with fresh PBS, avoiding disturbance of the gel surface. Samples were taken at: 2, 6, 9, 13, 23, 29, 36, 48, 54, 72 hrs and 4, 5, 7, 9, 11, 14, 18, 21, 25, 32, 39 days.

For dialysis release, PNP hydrogel loaded with bimatoprost (100 μL, 0.25 mg mL^−1^) was injected into mini dialysis tubes (molecular weight cutoff of 12 – 14 kDa; Sigma-Aldrich Pur-A-Lyzer). Each tube (n = 3) was sealed with parafilm, placed in PBS (5 mL) in a sealed 15 mL Falcon tube, and stored upright at 37 °C. At each time point, 250 μL of PBS was removed and replaced with an equal volume of fresh PBS. Samples were taken at: 2, 6, 9, 13, 23, 29, 36, 48, 54, 72 hrs and 4, 5, 7, 9, 11 days.

At the end of the study, gel was collected from each capillary or dialysis tube, diluted with PBS and remaining bimatoprost was quantified. Bimatoprost concentration in releasate was quantified using HPLC-MS as described in supplemental methods. Data is presented as bimatoprost remaining in gel, calculated as 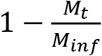, where *M_t_* is the amount released at each time point and *M_inf_* is the total amount loaded in the gel at the beginning of the assay. Data were fit with a one phase-decay in GraphPad Prism and the half-life of release was determined. Ritger-Peppas analysis was performed by fitting data with power law 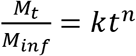 in GraphPad Prism to find *k* and *n*.

### In vitro dynamic Apparatus 7 release assay

Dynamic release was performed using the 400-DS Apparatus 7 instrument (Agilent Technologies, Santa Clara, CA) at 37 °C in a 10 mL cell. Each reciprocating sample holder (n = 2) contained 100 μL of PNP hydrogel with bimatoprost (0.74 mg mL^−1^) and the dissolution media was PBS (5 mL), with one dipping cycle per minute (1 DPM). At each sampling point, 0.1 mL of the medium was withdrawn through the auto-sampling port into an HPLC vial. The dissolution duration was 72 hours and samples were taken at: 0.5, 4, 8, 24, 36, 48, 60, and 72 hrs. Release profiles for bimatoprost and hydrogel component HPMC-C12 and PEG-PLA NPs were quantified using HPLC-CAD as described in supplemental methods. Data were fit with a one phase-decay in GraphPad Prism and the half-life of release was determined. Ritger-Peppas analysis was performed by fitting data with power law 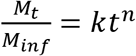 in GraphPad Prism to find *k* and *n*.

### Intravitreal administration of PNP hydrogel in rabbits

All animal procedures were conducted in accordance and compliance with internal IACUC approved standards and the ARVO *Statement for the Use of Animals in Ophthalmic and Vision Research*. Eight male New Zealand white (NZW) rabbits (~6 months old) received a single bilateral dose of 50 μL PNP hydrogel alone or with bimatoprost (0.16 mg mL^−1^) by ITV injection as described in **Table S1**.

Rabbits were placed under general anesthesia by intramuscular (IM) cocktail of 7 mg kg^−1^ ketamine HCl, 0.005 mg kg^−1^ dexmedetomidine HCl, and 0.1 mg kg^−1^ hydromorphone. Once appropriately sedated, a maintenance dose of isoflurane was delivered through a V-gel tube. To induce pupil dilation, 1% tropicamide was applied to both eyes prior to dosing procedure. A wire speculum was inserted. The eyelid margins and conjunctiva overlying the injection site were sterilized with 5% ophthalmic povidone iodine solution and 0.5% proparacaine hydrochloride was administered as a topical anesthetic. PNP hydrogel was administered via ITV injection into the inferotemporal quadrant of the globe using a 31-gauge x 8 mm needle, affixed to an insulin syringe. Upon completion of hydrogel administration to both eyes, isoflurane administration was halted, v-gel tube removed, and atipamezole administered for reversal of dexmedetomidine HCl.

### Ophthalmic Examination and Imaging

Ophthalmic examinations (OE), including slit-lamp biomicroscopy, indirect ophthalmoscopy, and rebound tonometry, were conducted to assess any intraocular changes and to characterize the hydrogel for up to eight weeks following ITV injection.

Spectral Domain Optical Coherence Tomography (SD-OCT) imaging occurred sequentially on sedated rabbits using a Heidelberg Engineering Spectralis with attached 30-degree lens to document any retinal changes. SD-OCT imaging captured several scans including a radially oriented line scan running underneath the injected PNP hydrogel. Wide-field color fundus and VH imaging (RetCam 3 with attached 130-degree lens) and B-scan ultrasonography (Philips Epiq 7G) documented the visibility and position of the injected PNP hydrogel.

### Necropsy with histopathology and PK analysis

Following euthanasia, eyes were enucleated and dissected. For eyes designated for histopathology, the globe with optic nerve was fixed in modified Davidson’s fixative for approximately 48 hours, then transferred to 70% ethanol for histologic evaluation. Histologic evaluation was completed by StageBio (Mason, Ohio). An axial superoinferior section was taken to include the optic nerve head and lens, with 3 step sections taken at 100 μm intervals. The temporal and nasal calottes were placed axial-side down and 3 step sections were taken at 100 μm each following embedding in paraffin. Sectioned blocks were stained with hematoxylin and eosin (H&E). H&E stained tissues were evaluated by an ACVP-board certified veterinary pathologist.

For eyes designated for PK analysis, VH was harvested and frozen at −70 °C. Prior to analysis, material was thawed and 100 μL of each VH sample was pipetted into individual cluster tubes, vortexed for 10 minutes, and centrifuged at 3700 rpm for 10 minutes at 4 °C. After centrifugation, supernatant was directly injected onto the autosampler for LC-MS/MS detection as described below.

### Bioanalytical method for bimatoprost PK results

A Nexera UPLC system (Shimadzu, Kyoto, Japan) with a Phenomenex XB-C18 column (50 × 2.1 mm, 2.7 μm) was used to analyze bimatoprost in VH. The gradient elution was 0.1% formic acid in water and 0.1% formic acid in acetonitrile with flow rate 1.1 mL min^−1^. A QTrap®5500 tandem mass spectrometer (Sciex, Foster City, CA) with Turboionspray (TIS) interface was operated in positive ionization mode with multiple reaction monitoring (MRM) for LC-MS/MS analysis. Bimatoprost standard was purchased from BioVision Inc and prepared at 1 mg mL^−1^ in 100% DMSO for the stock solution and serially diluted for a calibration curve from 2000 ng mL^−1^ to 0.1 ng mL^−1^.

## Acknowledgements

E.L.M. and R.A. contributed equally to this work. E.L.M. and E.A.A. conceived of the idea. E.L.M. performed experiments. Other authors aided with experiments. E.L.M. was supported by the NIH Biotechnology Training Program (T32 GM008412). C.M.K. was supported by the Stanford Bio-X William and Lynda Steere Fellowship. A.K.G. is appreciative of a National Science Foundation Graduate Research Fellowship and the Gabilan Fellowship of the Stanford Graduate Fellowship in Science and Engineering. A.l.d. was supported by the Schmidt Science Fellows program, in partnership with the Rhodes Trust. We thank Vladimir Bantseev, Donna Lee, Justin Ly, Liling Liu, and Melissa Schutten, current and former Genentech, Inc employees, for their scientific support and expertise.

