## Supplemental Materials for "Injectable polymer-nanoparticle hydrogel for the sustained intravitreal delivery of bimatoprost"

### 1. Supplementary Methods

*Vitreous humor mimic:* Vitreous humor mimic was created following literature reported protocol.[1] Briefly, sodium hyaluronate (1.01 MDa - 1.8 MDa; 28 mg) and agar (38 mg) were added to PBS (40 mL) and heated to boiling with stirring. Once fully dissolved and solution boiled, cuvettes were filled and allowed to set overnight prior to 50 μL gel and PBS injections.

*Bioanalytical method for bimatoprost static release assays:* An Agilent 1260 Infinity Binary HPLC and Agilent 6420 Triple Quadrupole mass spectrometer (Agilent Technologies, Santa Clara, CA) with a Zorbax EclipsePlus C18 column (2.1 x 50 mm, 1.8 μm; Agilent Technologies) was used to analyze bimatoprost in static release assay samples. The gradient elution was 0.1% formic acid in water and 0.1% formic acid in acetonitrile, operating with electrospray ionization in positive ion mode, source gas temperature 350 °C, gas flow rate 11 $min^{-1}$, and nebulizer pressure 40 psi. Data collection was performed using MassHunter Workstation LC/MS Data Acquisition software (Agilent) and bimatoprost was identified and quantified by integrated peak area in MassHunter Workstation Qualitative Analysis Navigator software (Agilent). Bimatoprost calibration curve was prepared in MilliQ water from 0.01 μM to 10 μM.

*PNP hydrogel characterization by High Performance Liquid Chromatography Coupled with Charge Aerosol Detector:* The PNP hydrogel composition was characterized by an HPLC system coupled with a charge aerosol detector (HPLC-CAD). The HPLC system used was Agilent 1260 series (Agilent Technologies, Santa Clara, CA, USA) equipped with a quaternary pump, vacuum degasser, temperature controlled autosampler, thermostatted column compartment, diode array detector, and coupled to a Thermo Dionex Corona Veo RS CAD detector (Thermo Fisher Scientific, Waltham, MA). Empower (Waters, Milford, MA) was used for data analysis.

*Endotoxin Measurement:* Endotoxin level in the PNP hydrogels were assessed prior to in vivo dosing using a kinetic Limulus Amebocyte Lysate (LAL) assay (Charles River Laboratories) with a 4-point standard curve (0.005 to 5.0 EU $mL^{-1}$). Control standard endotoxin (i.e., E.coli 055:B5 lyophilized) was reconstituted to a stock concentration of 50 endotoxin units (EU) $mL^{-1}$ in USP grade sterile water for irrigation (WFI). The PNP hydrogel was dissolved in 10% DMSO with a serial dilution in WFI. Data were analyzed using WinKQCL software.

### 2. Supplementary Data

| 2-month ITV Study Design in NZW Rabbit | | | | | | | |
|---|---|---|---|---|---|---|---|
| Group | Number of Animals | Treatment Group | Hydrogel Dose ($\mu L \cdot eye^{-1}$) | Bimatoprost Dose ($\mu g \cdot eye^{-1}$) | Route of Administration | Necropsy Study Day | PK Necropsy Study Day |
| 1 | 2 Males | PNP Hydrogel | 50 | 0 | ITV injection (bilateral) | 58 (n = 2) | N/A |
| 2 | 6 Males | PNP Hydrogel + Bimatoprost | 50 | 8 | Intravitreal (bilateral) | 58 (n = 2) | 29 (n = 2)<br>58 (n = 2) |

**Table S1**: The tolerability study design in NZW rabbits; Day 1 is the day of intravitreal administration of treatment.

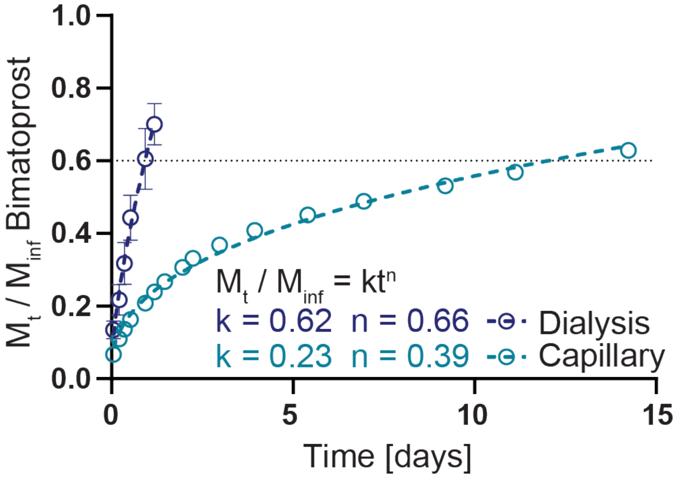


**Figure S1** – Ritger-Peppas fit of static release assay. 100 µL of PNP hydrogel with bimatoprost cargo (0.25 mg $mL^{-1}$) was injected into the bottom of a capillary or dialysis tube and PBS buffer added. Buffer was sampled over time to quantify bimatoprost release by LC-MS. Data are shown as mean ± SD (n = 3) and fit with the power law $\frac{M_t}{M_{inf}} = kt^n$ in GraphPad Prism and values for release constant ($k$) and diffusional exponent ($n$) calculated.

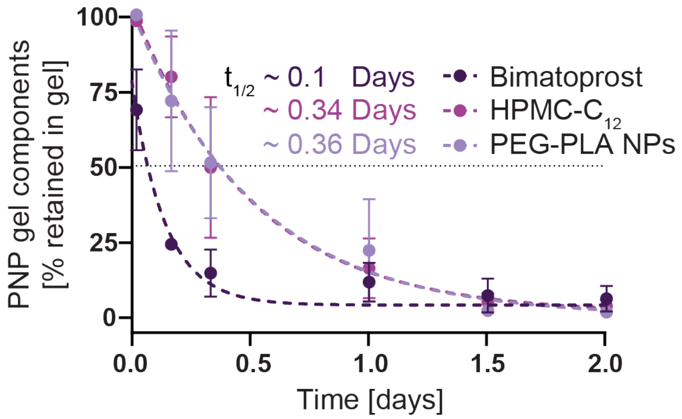


**Figure S2** – 400-DS Apparatus 7 release profile for PNP hydrogel component polymers and loaded bimatoprost. 100 µL of PNP hydrogel with bimatoprost (0.25 mg $mL^{-1}$) was injected into the mesh basket of the USP7 instrument, PBS buffer added and sampled over time while the basket was dipped at 1 cycle per minute. Release of bimatoprost and PNP hydrogel component polymers HPMC-$C_{12}$ and PEG-PLA was determined by RP-CAD. Data are shown as mean ± SD (n = 2) and fit with a one-phase exponential decays in GraphPad Prism and half-lives calculated.

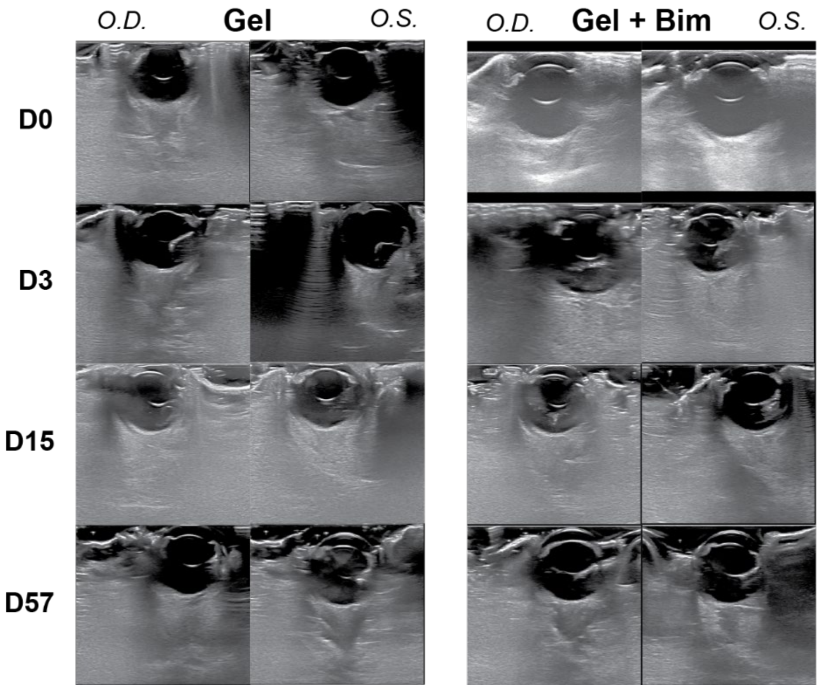


**Figure S3** – Representative ultrasound images from Gel and Gel + Bim treated NZW rabbit eyes. Images demonstrate subjectively no retinal changes over time after ITV injection of PNP hydrogel with or without bimatoprost. *O.D.* = oculus dexter, *O.S.* = oculus sinister.
